# Tracking single neurons across days reveals acute and persistent effects of psilocybin in the mouse brain

**DOI:** 10.64898/2026.11.14.766632

**Authors:** T.E. DeWeese, A. Cheng, S. Koukuntla, K.L. Tamashiro, T.D. Harris, A.R. Graves

## Abstract

The psychedelic psilocybin offers a novel approach to treat depression and other psychiatric disorders. A single dose of psilocybin induces rapid neuroplastic effects that persist for many weeks to months. However, psilocybin’s effect on neural firing beyond its systemic presence, and the neural mechanisms linking these acute and sustained effects remain unknown. Using chronically implanted Neuropixels probes, we tracked 1,692 neurons spanning prefrontal and hippocampal regions following saline and psilocybin injections and 24 h later. Psilocybin bidirectionally modulated firing acutely in a cell-type and region-specific manner. Persistent firing (24 h post-injection) is net suppressed relative to saline, including in acutely activated neurons. The late acute response, but not early peak response, predicted a neuron’s firing 24 h later. To our knowledge, this is the first evidence that directly reveals psilocybin’s sustained impacts on neuronal firing, providing key insight into the circuitry accompanying its lasting cognitive and behavioral alterations.

## Introduction

Psychedelics, particularly psilocybin, have received significant attention as a novel treatment for psychiatric disorders^1–3^. After one or two doses, psilocybin provides rapid and long-lasting antidepressant and anxiolytic effects in patients that persist for months after exposure^4–6^. These clinical results have been replicated in rodent models, with multiple studies observing decreased anxiety- and depression-like behaviors^7–10^, enhanced social processing^11,12^, and increased cognitive and behavioral flexibility^13^ following psychedelic exposure. As these behavioral benefits outlast the drug’s metabolites, a central question is what neural alterations persist after psilocybin is cleared from circulation.

Psilocybin elicits varied and conflicting neural responses across the brain, likely reflecting its complex pharmacology: it agonizes receptors with opposing effects (e.g., excitatory 5-HT_2A_ vs inhibitory 5-HT_1A_) and acts on both excitatory and inhibitory neural populations^14^. These receptors couple to distinct signal transduction pathways, such as Gq and β-arrestin signaling^15^, which can in turn trigger receptor internalization^15^, shifts in the AMPA/NMDA ratio^8^, and increased synaptogenesis^14^. Together, these processes unfold over distinct timescales. In mice, the acute response begins within minutes, as indexed by head-twitch response (HTR), and peaks 10 min after a 1 mg/kg subcutaneous dose^16^ and reflects 5-HT_2A_ activation^8,17^. Within hours, immediate early gene expression rises^18^, and within 24 h dendritic spine density increases in frontal cortex and remains elevated for weeks^7^.

Beyond these receptor-level and plasticity effects, psilocybin also alters activity at the systems level, with previous research identifying changes in brain regions associated with sensory processing, attention, memory, mood regulation, and brain-wide association, including the sensory and motor cortices, prefrontal cortex, hippocampus (HPC; particularly ventral HPC [vHPC]), and thalamus^19–25^. However, many recording methods used in these studies (e.g., fMRI, PET) indirectly infer neural activity from hemodynamic or metabolic signals, which are confounded by serotonin receptor–mediated vasoregulation altering cerebral blood flow and neurovascular coupling^21,26^. These methods have limited spatiotemporal resolution and therefore cannot capture single-neuron spiking that is required to understand fundamental neural mechanisms.

High-density electrophysiology (e.g., Neuropixels) can isolate single-unit activity by directly measuring firing of neurons at sub-millisecond resolution across multiple recording sites. Combined with histology, recorded units can be assigned to brain regions^27^, and waveform and firing properties can illuminate putative cell-type classes^28^. Recent single-unit studies in rodents have begun to resolve how psilocybin acutely alters neural activity, reporting heterogeneous effects. In the medial prefrontal cortex (mPFC), psilocybin elevated firing rates in pyramidal tract neurons but not in intratelencephalic neurons^29^. In GABAergic neurons, psilocybin reduced firing in somatostatin-expressing (SST) interneurons, increased activity in parvalbumin-expressing (PV) interneurons, and did not affect vasoactive intestinal peptide-expressing (VIP) interneurons^30^. In contrast, another study found that psilocybin modestly reduced regular-spiking neuron firing in the cortex, while narrow neurons (i.e., putative PV interneurons) were unaltered^31^. In retrosplenial cortex, psilocybin elevated spiking activity of mPFC-projecting neurons^32^. Psilocybin reconfigured burst coding rather than mean firing rate across hippocampal and thalamic circuits, and reduced firing rates in deep layers of mPFC^31^. A chronic Neuropixels study in rat mPFC found decreased net firing for about an hour after injection, and increases in beta and low-gamma local field potential (LFP) power that persisted for days, leaving persistent spiking effects unexplored^33^.

Together, these studies indicate that psilocybin alters firing rates acutely in a cell-type- and region-dependent manner. However, it remains unclear whether lasting changes in firing accompany the persistent structural, transcriptional, and behavioral effects, even though psilocybin’s enduring therapeutic benefits likely arise from persistent, network-level changes in neural activity. Addressing this requires chronic recordings that span both the acute and persistent phases. Chronically implanted Neuropixels probes can stably record hundreds of single units across distant brain regions for months^34^, and track individual units across days^35–38^. Yet these tools have not been used to track the same neurons from the acute drug effect to the persistent phase. Without tracking, a difference between sessions could reflect either a change in firing or a different sample of neurons. Following the same neurons would therefore reveal how their firing evolves from the acute drug effect to the persistent effect.

Here, we used up to 2 chronically implanted Neuropixels 2.0 probes in each of 8 head-fixed mice to record neural activity from 1692 single units across post hoc identified brain regions. We collected recordings immediately and 24 h following subcutaneous (s.c.) injection of saline and psilocybin, a time when structural plasticity is already detectable^7^. We targeted mPFC and vHPC because they are interconnected circuit nodes that regulate mood and are known to be altered by psilocybin^39^, and we also sampled thalamic nuclei along the vHPC probe track. We found psilocybin bidirectionally modulated firing in a cell-type- and region-specific manner, with the strongest increase in narrow putative PV interneurons in mPFC and vHPC, and no significant acute change compared to saline in the thalamic nuclei. Twenty-four hours later, there was a net suppression of firing, including among those units that had been acutely activated, and each unit’s acute response at 55-60 min, a single half-life^40^, was associated with its persistent outcome. These results reveal that the same neurons can shift from acute activation to persistent suppression, and provide a single-neuron evaluation of firing following a single dose of psilocybin that relates acute drug response to lasting changes in prefrontal and hippocampal neuronal activity.

## Results

### Chronic Neuropixels recordings tracked single neurons across saline and psilocybin sessions

To track neural activity across the acute and persistent phases of psilocybin’s effects, we implanted two chronic Neuropixels 2.0 probes in adult male mice targeting the medial prefrontal cortex (mPFC), ventral hippocampus (vHPC), and thalamic nuclei. Acute (immediately after injection) and chronic (24 h later) recordings were made from head-fixed mice viewing a 0.5-Hz sinusoidal grating to maintain consistent visual input across sessions (Fig. 1A–C). After handling and habituation to head-fixation, each mouse received subcutaneous saline injections on three baseline days (days −3 to −1), and psilocybin (1 mg/kg, s.c.) on Day 0, with recordings spanning 15 min before and 15 min (saline) or 60 min (psilocybin) after injection (Fig. 1A). We then collected additional 15 min recordings 24 h later (Day 1), when psilocybin is cleared from circulation.

**Figure 1.**
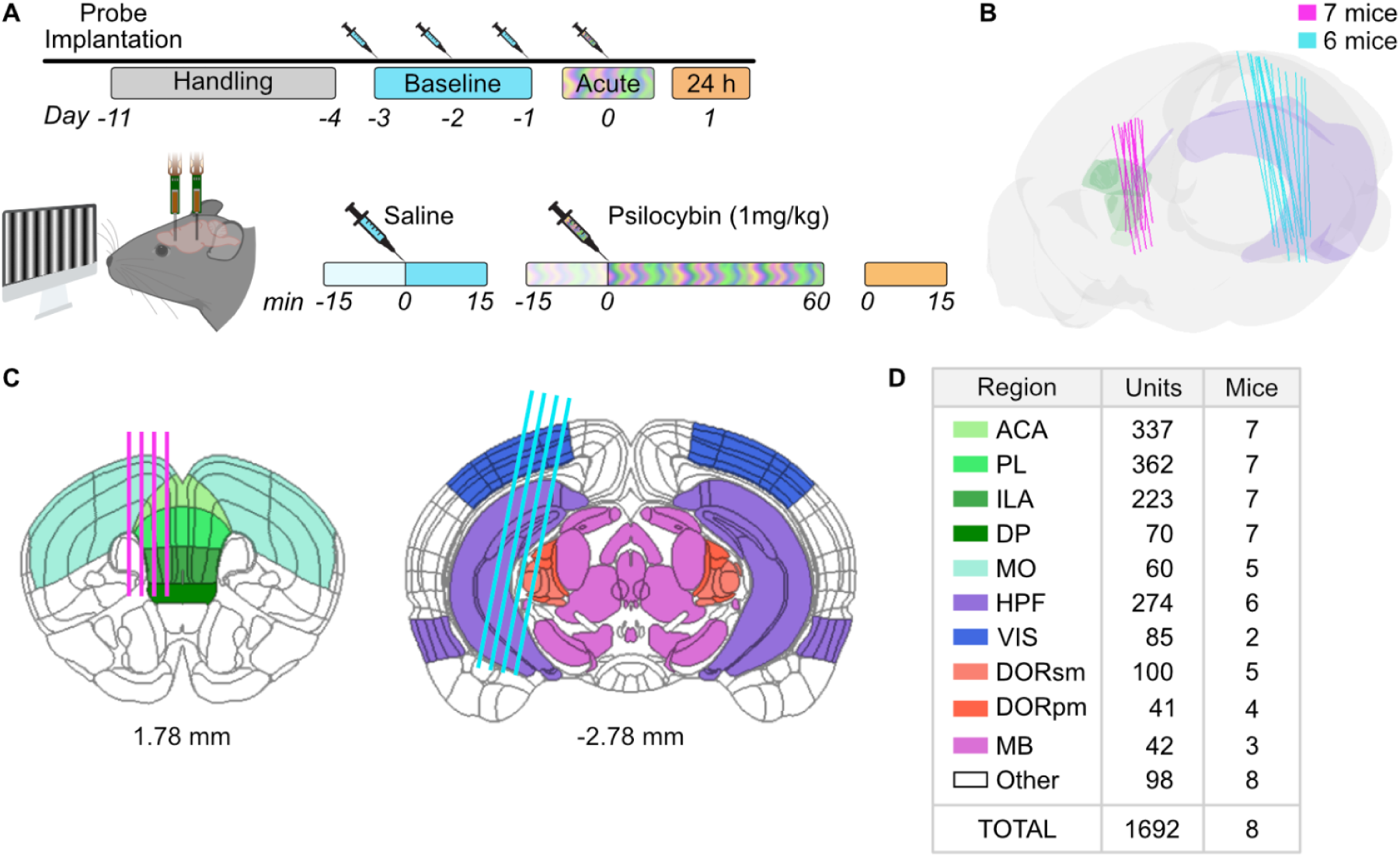
Chronic multi-region Neuropixels recording in head-fixed mice. **(A)** Chronic recording timeline. Mice were handled and habituated to head-fixation for 7 days (−11 to −4). On baseline days −3 to −1, mice received saline injection (s.c.) and were recorded for 30 min. On Day 0 (acute), mice received psilocybin (1 mg/kg, s.c.) and were recorded for 15 min before and for 60 min after injection. Additional 15 min head-fixed recording was collected 24 h (Day 1) after psilocybin (Day 0). A 0.5 Hz sinusoidal grating was displayed during all recordings. **(B)** 3D rendering of reconstructed Neuropixels probe tracks in medial prefrontal cortex (mPFC) (magenta, n = 7 mice) and ventral hippocampus (vHPC) (cyan, n = 6 mice) in the Allen CCFv3 brain (side view, anterior to the left). Only shanks with active recording sites are rendered. Tracks for the 2 mPFC mice and 1 vHPC mouse without histology are rendered from the known implantation trajectories. In total, 8 mice contributed to recordings. **(C)** Coronal sections at +1.78 mm and −2.78 mm relative to bregma, showing targeted probe placement (magenta, mPFC; cyan, vHPC). Regions are colored as in D. **(D)** Number of well-isolated units from each region tracked across the last baseline recording (Day −1), acute day (Day 0), and chronic Day 1 (Day 1). *ACA: anterior cingulate area, PL: prelimbic area, ILA: infralimbic area, DP: dorsal peduncular cortex, MO: somatomotor areas, HPF: hippocampal formation, VIS: visual areas, DORsm: sensory-motor associated thalamic nuclei, DORpm: polymodal association thalamic nuclei, MB: midbrain*

Due to the sustained transcriptional, structural, and network effects of psilocybin^7,14,41^, psilocybin injection always followed saline injection to minimize any baseline measurement contamination. In addition, to further stabilize baseline measurements, the final saline and psilocybin sessions were both performed in animals habituated with two saline injection sessions. The final saline session therefore served as a within-animal control for the non-specific effects of injection, head-fixation, and day-to-day variability in firing. This control is necessary because head-fixation may cause a stress response in animals and raise corticosterone, which can alter neural firing^42^.

Across 8 mice, we analyzed 7 mPFC probes and 6 vHPC after excluding damaged probes (Fig. 1B, C). The mPFC probes sampled the anterior cingulate (ACA, n=7), prelimbic (PL, n=7), infralimbic (ILA, n=7), and dorsal peduncular (DP, n=7) areas, as well as somatomotor cortex (MO, n=5). The vHPC probes sampled the hippocampal formation (HPF, n=6), visual cortex (VIS, n=2), sensory-motor and polymodal thalamic nuclei (DORsm, n=5; DORpm n=4, respectively), and midbrain (MB, n=3) (Fig. 1D) (see Methods for more narrowed region directions).

To track single-neuron activity across days, we concatenated each mouse’s Day −1, Day 0, and Day 1 recordings, preprocessed and spike-sorted the resulting recording, and selected well-isolated units present in all three sessions (Methods). This process yielded 1692 tracked units across 8 mice (Fig. 1D). Across days, 98% of units best matched their own waveform from a different day rather than that of any neighboring unit. Classifying each unit by its mean waveform and autocorrelogram shape yielded 334 narrow interneurons, 383 wide interneurons, 881 pyramidal cells, and 94 non-somatic units, whose waveforms are consistent with an axonal or dendritic rather than somatic spikes (Methods)^28^.

### Acute psilocybin bidirectionally modulated firing in a cell-type- and region-specific manner

We first examined how acute psilocybin changes the firing of tracked neurons. Psilocybin (1 mg/kg, s.c.) is known to evoke a head-twitch response in mice that peaks 10 min after injection^16^. We focused on the time window 10–15 min post-injection, which coincides with this peak and the onset of psilocybin’s neural effects^16,29,32^. As units differ widely in baseline firing, we *z*-scored (see methods) the firing rate in this window against its own pre-injection baseline, thus representing firing as standard deviations from the baseline mean (see Methods). We classified units as activated (*z* > 2), suppressed (*z* < −2), or not modulated (|*z|* ≤ 2). After saline, 15.7% of units were classified as activated and 5.6% as suppressed; after psilocybin, 23.0% were activated and 13.7% were suppressed (n = 1692; Fig. 2A). The distribution of response classes differed significantly between conditions (p = 8.4 × 10⁻²⁶, Stuart-Maxwell), and psilocybin produced a larger fraction of both activated and suppressed units (activated: adjusted p = 6.3 × 10⁻⁸; suppressed: adjusted p = 6.9 × 10⁻¹⁷; McNemar’s exact, Bonferroni-corrected). Acute psilocybin therefore modulated firing in both directions rather than uniformly activating or suppressing the population. Because units from the same probe or subject are not independent, we confirmed all significant effects in this section with a Bayesian categorical mixed-effects model that accounts for subject, probe, and unit (Methods).

**Figure 2.**
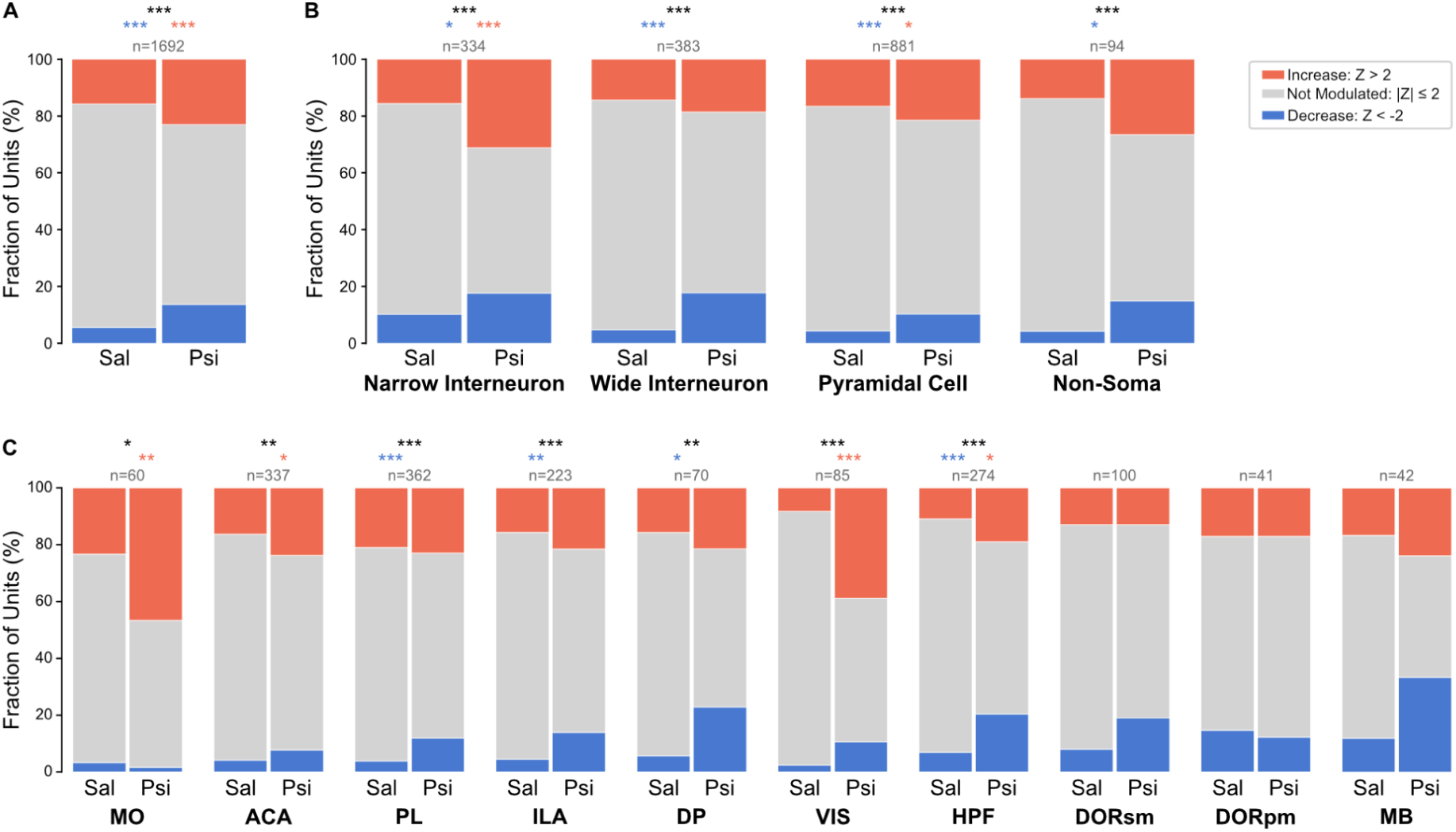
Acute psilocybin bidirectionally modulated firing across cell types and brain regions. **(A)** Acute firing-rate modulation by psilocybin (Psi, Day 0) compared with saline (Sal, Day −1). Only units tracked across both days included (n = 1692). For each unit, firing rate in 10–15 min post-injection window was *z*-scored to 10 min pre-injection baseline taken from center of 15 min pre-injection period (1 min bins). Units with *z* > 2 were classified as activated, units with *z* < −2 as suppressed, and all others (|*z*| ≤ 2) as not modulated. **(B)** Modulation by putative cell type (narrow interneuron, wide interneuron, pyramidal cell, and non-somatic units; see Methods). **(C)** Modulation by brain region. Statistical analyses: Stuart-Maxwell (black); Bonferroni-adjusted McNemar exact (red=activated, blue=suppressed). A Bayesian categorical mixed-effects model with random intercepts for subject, probe, and unit confirmed all significant effects as credible (95% CrIs excluding 1; Methods). Asterisks: *p < 0.05, **p < 0.01, ***p < 0.001

This bidirectional modulation was cell-type dependent (Fig. 2B): all four putative cell types (classified with CellExplorer^28^; Methods) had a significant shift in response between saline and psilocybin (all p < 0.001, Stuart-Maxwell). A larger fraction of narrow interneurons, which are thought to include parvalbumin-expressing cells, was activated after psilocybin than after saline (15.6% saline vs 31.3% psilocybin; p = 1.9 × 10.⁻⁶; McNemar exact, Bonferroni-corrected), the clearest activation of any cell type. A modestly larger fraction was also suppressed (10.2% vs 17.7%; p = 0.010). Wide interneurons had the largest difference in suppressed units between conditions (4.7% vs 17.8%; p = 1.0 × 10.⁻⁸) but no significant difference in activated units (14.4% vs 18.5%; p = 0.27). In putative pyramidal cells, psilocybin increased the proportion of both suppressed (4.3% vs 10.3%; p = 1.3 × 10.⁻⁶.) and activated units (16.6% vs 21.5%; p = 0.016). In non-somatic units, whose waveforms suggest axonal or dendritic signals, suppressed units were also more frequent after psilocybin (4.3% vs 14.9%; p = 0.013), whereas the difference in activated units did not reach significance after correction (13.8% vs 26.6%, p = 0.071). Thus, psilocybin increased the percentage of suppressed units in every cell type, whereas activated units were more frequent only in narrow interneurons and, to a lesser extent, pyramidal cells.

The modulation also varied across regions (Fig. 2C). Psilocybin significantly shifted response classes in every region except the thalamic nuclei and midbrain (Stuart-Maxwell, p < 0.05 in all other regions; DORsm: n = 100, p = 0.10; DORpm: n = 41, p = 0.94, MB: p = 0.050). Within the mPFC, the shift differed by dorsal/ventral subregion. Suppressed units were more frequent after psilocybin in ventral regions (PL, ILA, DP), most significantly in PL (PL: 3.9% saline vs 11.9% psilocybin; p = 5.0 × 10.⁻⁵; ILA: p = 0.0010; DP: p = 0.015; McNemar’s exact, Bonferroni-corrected), whereas activated units were more frequent in dorsal regions (ACA: 16.3% vs 23.7%, p = 0.023; MO: 23.3% vs 46.6%, p = 0.0087). Among regions sampled by vHPC probes, both activated units (p=0.021) and suppressed units (p = 3.9 × 10.⁻⁶) were more frequent after psilocybin in the HPF. While MB responses were not significantly different (p = 0.050), the increase in suppressed units (11.9% vs 33.3%) was credible in the mixed-effects model relative to saline (OR = 6.6, 95% CrI 1.8–29.0, pd = 0.999). VIS had the largest difference in activated units (8.2% vs 38.8%; p = 5.1 × 10.⁻⁶) and was credible in the mixed-effects model that accounts for subject and probe (OR = 13.3, 95% CrI 4.1–49.9, pd > 0.999), although these units came from only 2 mice and should be interpreted with caution.

### Psilocybin produced persistent net suppression of firing 24 h later

We next examined whether these changes persisted 24 h after psilocybin injection. Using the same 1692 units, we quantified firing 5–10 min into the recording 24 h after psilocybin (Day 1), and computed each unit’s firing rate *z*-score 24 h after injection relative to the same pre-injection baseline as in Fig. 2 (Fig. 3A). We compared this with the same measurement 24 h after the last saline injection, taken from the Day 0 pre-injection period. Saline sessions had substantial day-to-day variability in single-unit firing, with 25.7% of units classified as activated and 16.1% as suppressed. Even so, the response-class distribution 24 h after psilocybin differed significantly from saline (p = 1.9 × 10.⁻¹¹, Stuart-Maxwell): the fraction of activated units fell to 18.7% (vs 25.7%; p = 6.6 × 10.⁻⁶; Bonferroni-adjusted McNemar’s exact) and the fraction of suppressed units rose to 25.6% (vs 16.1% after saline; p = 2.9 × 10.⁻¹¹). A Bayesian mixed-effects model confirmed both effects (suppressed: OR = 1.73, 95% CrI 1.45–2.09, pd > 0.999; activated: OR = 0.77, 95% CrI 0.65–0.92, pd = 0.997).

**Figure 3.**
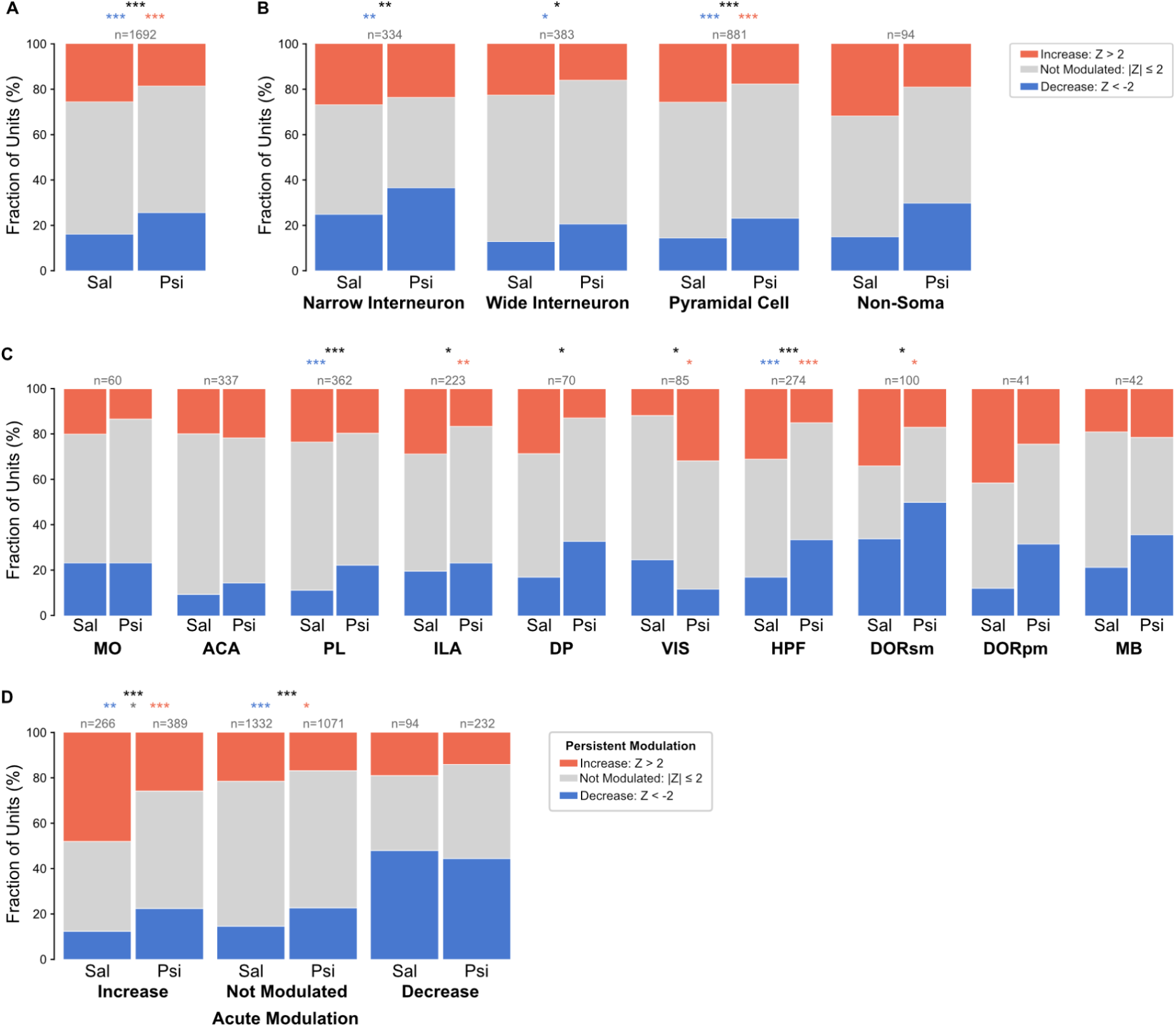
Psilocybin produces persistent, cell type- and region-specific shifts in firing 24 h after administration. (A) Persistent firing-rate modulation 24 h after psilocybin (Psi; chronic Day 1, compared with 24 h after saline (Sal; Day 0 pre-injection period, 24 h after the last saline injection)). The same 1692 tracked units as in Fig. 2 were analyzed. Units were classified as activated (*z* > 2), suppressed (*z* < −2), or not modulated (|*z*| ≤ 2) as in Fig. 2 when measured 5–10 min into the 24 h recording. (B) 24 h modulation by putative cell type. (C) 24 h modulation by brain region. (D) Persistent (24 h) response classes as a function of acute response for saline and psilocybin. Units are grouped by acute response (x-axis: increase = activated, not modulated, decrease = suppressed; n above each bar). Stacked segments show the fraction of those acute units in each persistent class. Statistical analysis: (A-C) Stuart-Maxwell test (black), Bonferonni-adjusted McNemar exact (red/blue); (D) χ2 contingency (black), Bonferonni-adjusted Fisher exact (red/gray/blue). A Bayesian categorical mixed-effects model with random intercepts for subject, probe, and unit confirmed all significant effects in panels A–C as credible (95% CrIs excluding 1), and three of the significant transitions in panel D (acute to persistent: increase to increase, increase to decrease, and not modulated to decrease; Methods). Asterisks: *p < 0.05, **p < 0.01, ***p < 0.001

The shift toward suppressed firing 24 h post psilocybin was present across all four putative cell types (Fig. 3B). Response classes shifted significantly in each cell type (p < 0.05; Stuart-Maxwell), except non-somatic (p = 0.057). Within these cell types with significant shifts, the fraction of suppressed units was elevated in each (all p < 0.05; McNemar’s exact, Bonferroni-corrected). Only pyramidal cells also had significantly fewer activated units relative to saline (25.8% saline vs 17.8% psilocybin; p = 0.00029). Regionally, persistent shifts were significant in PL, ILA, DP, HPF, VIS, and DORsm, but not in ACA, MO, DORpm, or MB (Fig. 3C). PL and HPF had more suppressed units (PL: 11.3% vs 22.4%, p = 0.00016; HPF: 17.2% vs 33.6%, p = 3.2 × 10.⁻⁵), and ILA, HPF, and DORsm had fewer activated units. VIS was the exception, with more activation (11.8% vs 31.8%; p = 0.012), although this came from only 2 mice. Within the mPFC, the ventral subregions that were acutely suppressed (PL, ILA, DP) also had persistent shifts (more suppressed units in PL, fewer activated units in ILA, and an overall shift in DP), whereas the dorsal subregions (ACA, MO) did not. A Bayesian mixed-effects model with random intercepts for subject, probe, and unit confirmed all of these significant effects (Methods).

To determine whether the same units were suppressed acutely and persistently, we mapped each unit’s acute response class onto its persistent class (Fig. 3D). Transitions differed between psilocybin and saline for acutely activated units (p = 5.1 × 10.⁻⁸ χ² test, Bonferroni-corrected) and acutely not-modulated units (p = 1.7 × 10.⁻⁶), but not for acutely suppressed units (p = 0.89). After psilocybin, acutely activated units were much less likely to remain activated and were more likely to return to baseline or become suppressed compared to saline (activation: 26.0% vs 48.1%, −22.2 percentage points, p = 4.35 × 10.⁻⁸; unmodulated: 51.7% vs 39.5%, +12.2%, p = 0.015; suppression: 22.4% vs 12.4%, +10.0%, p = 0.0082; Fisher exact, Bonferroni-corrected). Some units showed persistent modulation due to psilocybin despite not showing significant acute modulation: after psilocybin, 22.7% of units that were not acutely modulated became suppressed persistently (vs 14.6% after saline; +8.0%; p = 2.5 × 10.⁻⁶) and 17.0% became activated (vs 21.6%; −4.6%; p = 0.027). A Bayesian mixed-effects model, with not-modulated (n.s.) as the reference persistent class, found credible effects for three of the six transitions it can test. Acutely activated units were less likely to remain activated (OR = 0.45, 95% CrI 0.31–0.67, pd > 0.999) and more likely to become suppressed (OR = 2.42, 95% CrI 1.26–4.73, pd = 0.998). Acutely unmodulated units were also more likely to become suppressed (OR = 1.67, 95% CrI 1.34–2.13, pd > 0.999). The small shift of acutely not-modulated units toward activation was not credible in this model (OR = 0.84, 95% CrI 0.68–1.04, pd = 0.949). Acute psilocybin was therefore followed by a net shift toward suppression, including in a subset of acutely activated units (Fig. 3D).

### Time course of acute psilocybin response relates to its persistent modulation

To relate acute and chronic responses, we first examined the time course of firing in each acute response class (Fig. 4). After psilocybin, units classified as activated (n = 389, 23%) increased firing rate after injection to a mean z-score of +6.2 at 12.5 min, and suppressed units (n = 232, 14%) fell to −6.4 at 14.5 min, while not-modulated units (n = 1071, 63%) stayed near baseline (maximum deviation −0.26; Fig. 4A). Because classes are defined from the post-injection window, units classified as activated or suppressed after saline have similar ramps by construction, and they reach comparable *z*-scores (+6.2 and −4.9). Psilocybin nonetheless increased the magnitude of single-unit responses by 1.6x relative to saline (median |z| 1.42 vs 0.91; paired Wilcoxon signed-rank, p = 1.0 × 10^-34^; n = 1692 units, 8 animals; Fig. 4B, top). To determine whether this magnification reflected strong responses in a subset of units or a broad response pattern, we compared the full |z| distributions (Fig. 4B, bottom). Psilocybin |z| exceeded saline |z| at every decile (all 95% bootstrap CIs excluding zero), with a psilocybin/saline quantile ratio of 1.51–1.67. This steady increase (distance from identity) across the entire z-score distribution is consistent with psilocybin scaling response magnitude across the population, rather than recruiting a distinct subset of strongly responsive units.

**Figure 4.**
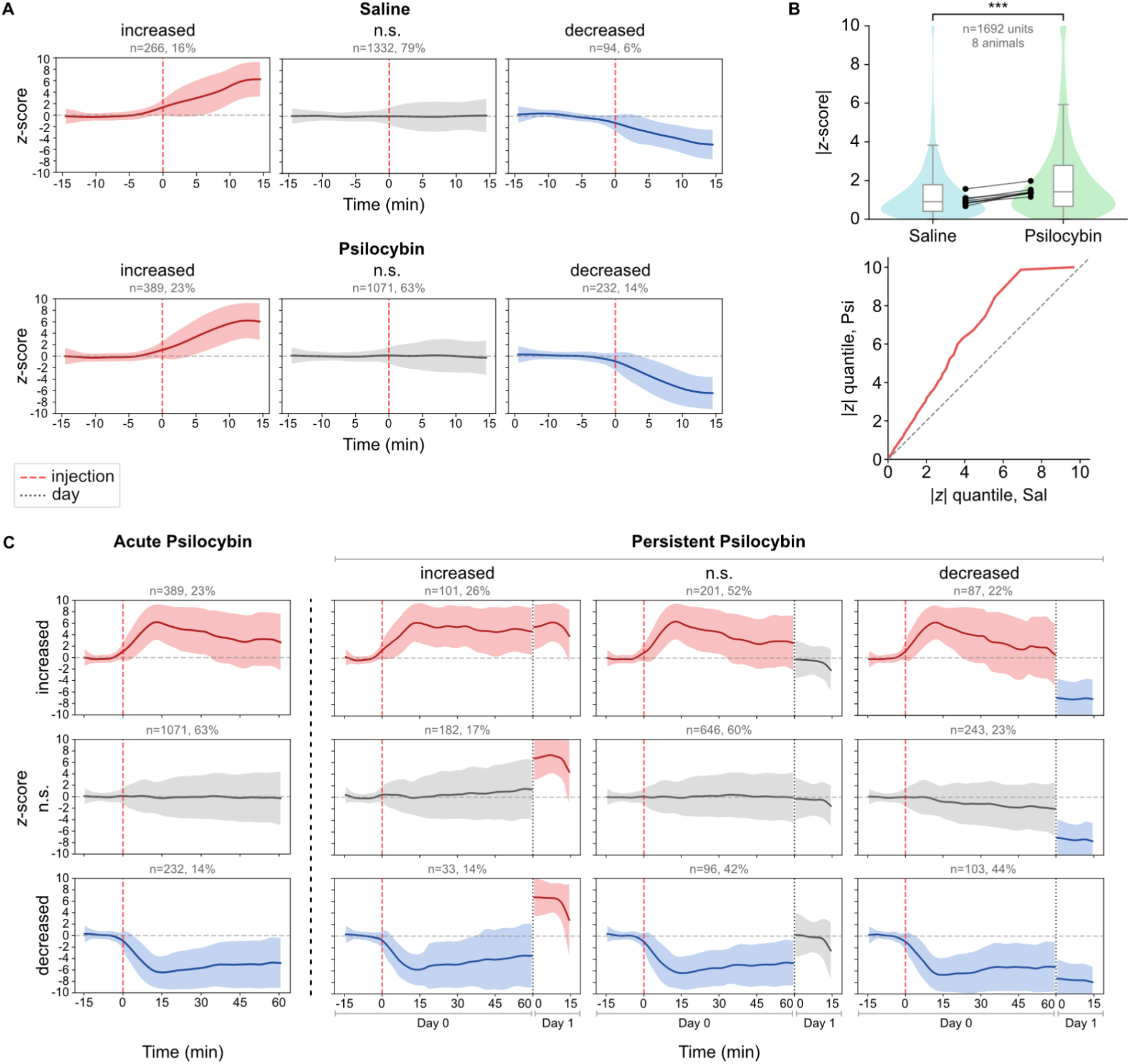
Time course of acute and persistent psilocybin modulation across response classes. **(A)** Mean firing-rate time course (*z*-score) of units grouped by acute response class (activated, not modulated [n.s.], suppressed) after saline (top row; Day −1) and psilocybin (bottom row; Day 0). Time is relative to injection (0 min; red dashed line). Psilocybin traces are truncated to the first 30 min of recording to match the saline sessions. Mean ± 1 s.d. across units is shown. n and percentages give the number and fraction of the 1692 tracked units in each class. The *z*-scores are relative to the pre-injection baseline (1 min bins, Gaussian-smoothed, sigma=2). **(B)** Top, magnitude of the z-scored firing-rate change (|z|) for each unit 10–15 min after injection, for saline (Day −1) and psilocybin (Day 0) (n = 1692 units, 8 mice). Each injection is z-scored against its own 10 min pre-injection baseline taken from the center of the 15 min pre-injection period, and z-scores were clipped to ±10. Violins show the distribution across units, with box plots inside (median and interquartile range; whiskers 1.5× IQR). Black dots show per-animal medians for each injection. The bracket shows a Wilcoxon signed-rank test on paired units. Bottom, quantile–quantile plot of |*z*| for psilocybin versus saline, with all units pooled within each condition. Curve plots the 1st–99th percentiles of |z| across units in psilocybin against the same percentile of saline. Dashed line: identity (no change). Plateau at |z| = 10 reflects z-score clipping. **(C)** Psilocybin time courses by acute and persistent response class. Left, full acute-day recording (Day 0; 75 min) for units grouped by acute response class (rows). Peak modulation occurs approximately 12–15 min after injection. Right, the same units are further divided by persistent response classes (columns; as in Fig. 3D), shown for the acute day (Day 0; −15 to 60 min relative to injection) and the chronic Day 1 recording (Day 1; 0–15 min), separated by the black dotted line. Day 1 *z*-scores are relative to the Day 0 pre-injection baseline. n and percentages are the number and row-wise fraction of units per acute class (as in Fig. 3D). Statistical analysis: (B) Wilcoxon signed-rank. Asterisks ***p < 0.001.

Over the full 75 min acute recording, peak modulation occurred 12–15 min after injection, close to the time at which the head-twitch response peaks at this dose^16^ (Fig. 4C, left). Dividing units by persistent class indicated that the acute trajectory was related to the persistent outcome (Fig. 4C, right). Among acutely activated units, those that stayed activated on Day 1 (n = 101) maintained a *z*-score of +5.1 with a slight dip to +4.79; those that returned to baseline on Day 1 (n = 201) relaxed to +2.8 by the end of the Day 0 recording, and those that became suppressed (n = 87) relaxed back to 1.29 by the end of the Day 0 recording. On Day 1, z-scored firing relative to Day 0 pre-injection in these units was +5.5, −0.65, and −7.11, respectively. Acutely suppressed units followed the same pattern: units that became activated on Day 1 (n = 33) had partially recovered by the end of the Day 0 to −3.4, whereas units that stayed suppressed (n = 103) remained at about −5.3. Acutely not-modulated units also drifted slightly upward (+1.4; n = 182) or downward (−1.9; n = 243) before becoming persistently activated or suppressed. To test this relationship across all units, we asked whether the late acute response (mean z-score, 55–60 min after injection) predicted Day 1 firing. In a linear mixed model with per-animal random slopes, the late acute response predicted Day 1 firing (β = 0.27, 95% CI 0.15–0.39, p < 0.0001; positive in 7 of 8 animals). When modeled with the early (13-18 min) mean z-score as an additional predictor, only the late response predicted Day 1 activity (β_early_ = -0.033, 95% CI -0.11–0.041, p = 0.38; β_late_ = 0.29, 95% CI 0.16–0.42, p < 0.0001). Together, these results suggest that a unit’s persistent outcome is related to how its acute response evolves over the first hour: units whose activation was sustained stayed activated, whereas units whose activation decayed ended up suppressed.

## Discussion

We found that psilocybin bidirectionally modulated firing relative to saline in a cell-type- and region-specific manner, both acutely and at 24 h. Acutely, psilocybin increased single-unit responses by about 1.6-fold across the population. Interestingly, we also found that a unit’s late acute response predicted the unit’s firing 24 h later. Because we tracked the same units following saline, acute psilocybin, and 24 h post-psilocybin sessions, differences between sessions reflect changes in firing of the same neurons, not changes in which neurons are sampled. Together, these results indicate that a single dose of psilocybin leaves a persistent, net suppressive change in prefrontal and hippocampal activity that, while indicated by the acute response, is not a simple prolongation of the acute response.

During drug exposure, suppression increased in every somatic cell type, most strongly in wide interneurons, a group that tends to include SST-expressing cells, whereas activation increased most commonly in narrow interneurons, a group that tends to include PV-expressing cells. Waveform width and autocorrelogram are only a proxy for cell identity; this pattern agrees with a recent optogenetic study that reports psilocybin increases PV interneuron activity and decreases SST interneuron activity^30^. Direct excitation of PV interneurons through 5-HT_2A_ receptors could drive the pyramidal suppression through perisomatic inhibition^43^, while SST interneurons can be inhibited directly through 5-HT_1A_ receptors^30^, which would fit the strong suppression of wide interneurons we observed.

Pyramidal cells had a significant increase in activated units alongside a proportionally larger increase in suppressed units. This pattern is consistent with psilocybin selectively exciting subpopulations such as subcortical-projecting PT neurons, while suppressing the larger intratelencephalic population^29^, although we could not separate these populations in the present study. Reduced SST activity could also disinhibit pyramidal dendrites and contribute to the activation, alongside direct 5-HT_2A_ excitation of apical dendrites^44^.

The acute effects of psilocybin varied across brain regions. Within mPFC, PL, ILA, and DP shifted toward suppression, whereas ACA and MO shifted toward activation. This may reflect differences between ventral and dorsal mPFC in 5-HT_2A_ expression, layer composition, or connectivity^45,46^. Ventral subregions that were acutely suppressed also displayed persistent shifts 24 h later, suggesting that activity-dependent plasticity mechanisms may engage acutely active neuronal ensembles to modify their chronic activity. However, units that were not acutely modulated also exhibited persistent firing changes, so acute modulation of a unit is not required, which may point to more than one plasticity mechanism. Broadly across the hippocampus, HPF had both more activated and more suppressed units, and MB had a non-significant but credible shift toward suppression, whereas the thalamic nuclei did not differ significantly from saline. These nuclei (DORsm: MG, PP, PoT; DORpm: LP, PIL, POL, SGN) are posterior thalamic nuclei and were sampled with relatively few units (DORsm, n = 100; DORpm, n = 41), so a small effect cannot be excluded. While our sampling omitted the midline and mediodorsal nuclei strongly linked to prefrontal-limbic mood circuits^47,48^, other DORsm or DORpm nuclei may also exhibit alterations to psilocybin. VIS had the largest increase in the fraction of activated units, consistent with psilocybin’s strong effects on sensory processing^49,50^. However, VIS was only captured from 2 mice and, although credible in the mixed-effects model, needs replication. Because the mice were viewing a video showing a sinusoidal grating along vertical axis during recording, this may reflect altered processing of visual input rather than a change in spontaneous activity, which we cannot distinguish without a no-stimulus condition.

Our acute results differ in part from a chronic Neuropixels study in rat mPFC, which found net suppression in both regular-spiking (corresponding to our pyramidal cells and wide interneurons) and narrow interneurons relative to saline^33^, whereas we found both suppression and activation. Several differences could contribute: species, route, and behavioral state, since those rats moved freely and alternated between rest and task blocks, while our mice were head-fixed and viewed a grating. Our results align with cell-type-resolved studies, which find activation and suppression in units based on cell type^29,30^. The chronic study also reported a decrease in broadband coherence and increases in beta and low-gamma LFP power, with LFP changes persisting for days. Purple et al. noted that gamma oscillations depend on PV interneuron activity^33,51^, these changes could reflect altered inhibitory dynamics involving PV cells or lasting neuroplastic changes, such as a shift in excitatory–inhibitory synaptic balance. In our data, a larger fraction of narrow-spiking interneurons was activated acutely, but 24 h later a larger fraction was suppressed. Pyramidal cells were also suppressed at 24 h. Because both interneurons and pyramidal cells were suppressed, the persistent change is not explained by a simple increase in inhibition. Instead, it may reflect a general reduction in network drive, or a lasting shift in excitatory–inhibitory balance. Unlike the chronic study, which reported persistence only in LFP, we also found persistent suppression of single-unit firing across cell types.

Twenty-four hours after psilocybin, firing was net suppressed rather than a prolongation of the acute effect. The transition analysis indicates that this suppression was not limited to neurons that were suppressed acutely. Relative to saline, which controls for day-to-day variability, acutely activated units were about half as likely to remain activated, and both acutely activated and not-modulated units were more likely to become suppressed. This net suppression may seem at odds with reports that a single dose of psilocybin increases dendritic spine density in mPFC for days to weeks^7^, but structural plasticity could accompany a reorganization of activity rather than a net increase in activity. Homeostatic mechanisms are known to scale down quantal amplitude globally to prevent runaway excitation when structural connectivity increases^52^, which may explain a delayed suppressive response. Although strong acute drive may trigger compensatory processes like long-term depression that scale down excitability afterward, the delayed suppression in units that were not modulated acutely points toward broader transcriptional shifts, such as altered immediate early gene expression and adjustment of the excitatory-inhibitory balance^18^. Though electrophysiological data alone cannot distinguish among these mechanisms, our findings suggest that mechanisms underlying the persistent effects of psilocybin must account for the loss of activation in acutely activated units and for suppression in units that were not acutely modulated.

The time course of the acute response was related to the persistent outcome. Among acutely activated units, those whose activation decayed back toward baseline by the end of the acute recording tended to be suppressed on Day 1, whereas units with sustained activation remained activated. Acutely suppressed units followed the same pattern, with partial recovery in the few units that became activated on Day 1. Across all units, the late acute response predicted Day 1 firing, and the early response added no additional predictive value. The predictive power of the late response may point to a process that depends on prolonged engagement of serotonin receptor signaling.

Persistent suppression of prefrontal and hippocampal firing may be relevant to the antidepressant effects of psilocybin. Hyperactivity in subgenual cingulate and related prefrontal circuits has been linked to treatment-resistant depression, and both clinical response to antidepressants and deep brain stimulation are associated with reduced activity in those regions^53,54^. In our data, ILA, the rodent homolog of subgenual cingulate^55^, along with PL and HPF, shifted toward suppression 24 h after psilocybin, a time at which antidepressant-like behavioral effects have been reported in mice^8^. This pattern is consistent with the idea that reduced activity in these circuits could contribute to psilocybin’s antidepressant effect. However, our mice were healthy and we did not use a stress model or measure behavior, so whether this suppression is related to psilocybin’s antidepressant-like effects remains untested. A stress model with a behavioral readout could test whether the degree of suppression in these regions predicts behavioral improvement.

Our study design has some limitations. Saline always preceded psilocybin because of psilocybin’s persistent effects. Therefore time, habituation, and recording drift cannot be fully separated from drug effects. However, we mitigated this by z-scoring each session to its own pre-injection baseline and tracking the same units across sessions, so each unit served as its own control. Injection itself also modulated firing after saline, likely reflecting injection stress during head-fixation. Subcutaneous injection provided a quicker, less-intrusive injection relative to an intraperitoneal route, but an indwelling catheter would likely reduce injection-related modulation further. We only used male mice (n=8), so sex differences could not be examined, however previous data have not reported firing differences between sexes^29–32,49^. Finally, findings cannot be correlated to attentional state, as the present study did not involve any behavioral recording during head-fixation.

Several experiments could extend these findings. Optogenetic tagging of PV, SST, and PT neurons would test the cell type mechanisms, and 5-HT_2A_ or 5-HT_1A_ antagonists would test which receptors are required for the acute and persistent changes, including whether preventing the acute response prevents a persistent one. Finally comparisons with other psychedelics and ketamine would show whether persistent suppression is a shared feature among rapid-acting antidepressants. Overall, our results link the acute pharmacology of psilocybin to lasting changes in prefrontal and hippocampal activity that may underlie its long-term behavioral effects.

## Methods

### Animals

Wild-type adult male C57BL/6J mice (8-15 weeks of age at the time of surgery; Jackson Laboratory) were used in this study. Male mice were used given their larger size to accommodate the weight of 2 probes and their fixtures. Animals were maintained on a 12:12 hr light/dark cycle with *ad libitum* access to food and water. After chronic probe implantation, mice were housed in semi-social pairs within partitioned rat cages separated by a transparent, perforated acrylic divider, permitting visual, auditory, and olfactory interaction while preventing physical contact and protecting hardware. The Institutional Animal Care and Use Committee (IACUC) at Johns Hopkins University approved animal care and experimental procedures.

### Surgical Procedures & Chronic Probe Implantation

Anesthesia was induced with 3% isoflurane in oxygen and maintained at 1-2% via nosecone on a stereotaxic frame (Kopf Instruments). Core body temperature was maintained at 37°C using a feedback-controlled heating pad. Extended-release buprenorphine (1.0 mg/kg, s.c.) was administered preoperatively for analgesia. The scalp was shaved, disinfected with alternating povidone-iodine and 70% ethanol scrubs, and infiltrated with 2% lidocaine hydrochloride prior to incision.

After the scalp incision, the skull was exposed, cleared of periosteum, and lightly scored to improve adhesive bonding. Bregma and lambda were leveled along the anteroposterior axes using a motorized stereotaxic manipulator (Sensapex). The skull surface was primed with light-curable adhesive resin (OptiBond Solo Plus, Kerr Dental). Slot craniotomy locations were marked using a high-speed dental drill, and a custom 3D-printed headplate was secured with UV-curable dental cement (RelyX Unicem 2, 3M). A stainless steel ground screw, pre-soldered to a platinum wire terminating in a gold connector pin, was inserted through a small craniotomy to contact the dura and anchored with dental cement.

Slot craniotomies were opened over the following coordinates relative to bregma: 1) mPFC: AP +1.79 mm, ML −0.20 mm, Depth −3.00 mm (vertical trajectory); 2) vHPC: AP −2.92 mm, ML −1.75 mm, Depth −5.31 mm (trajectory pitch angle: 9°). Exposed dura were irrigated with sterile saline to prevent desiccation.

Neuropixels 2.0 probes (4-shank, imec) were mounted in custom chronic fixtures and attached to a stereotaxic inserter (SCREWx^56^). Probes were lowered at 3–5 µm/s to target depths. Craniotomies were sealed with biocompatible silicone gel (Dura-Gel, Cambridge NeuroTech), and fixture assemblies were permanently anchored to the headplate and skull with dental cement before inserter removal. A single probe’s ground lead was connected to the ground screw via gold connector pins, with ground and reference tied. In one animal, the platinum ground wire broke, so the 4 shank-tip electrodes connected through shank switches were used as reference, rather than the external ground screw. The entire assembly was enclosed in a custom transparent acetate shroud.

### Electrophysiology Data Acquisition

Mice recovered for at least 48 h post-surgery and were habituated to head-fixation for 7 d before recording. Neural signals were acquired using Neuropixels 2.0 connected to a PXIe acquisition system (National Instruments) controlled by SpikeGLX software. Data were sampled at 30 kHz.

### Electrophysiological Data Preprocessing

Raw data were preprocessed using CatGT^57^, including phase-shift correction for multiplexed sampling, bandpass filtering (300–10,000 Hz, Butterworth), artifact removal, and global median referencing. Recordings spanning multiple experimental sessions were concatenated into unified binary files (Supercat^57^). Non-rigid motion correction was performed using DREDge with default parameters^58^.

Spike sorting was performed using Kilosort4^59^ with intrinsic drift correction disabled to preserve DREDge motion correction. To address cluster oversplitting, initial clusters meeting baseline quality thresholds (SNR > 2.0, total spike count > 100, and refractory period contamination < 0.1) were processed using the SLAy^60^ auto-merging framework.

### Unit Classification and Quality Metrics

Automated unit curation followed criteria adapted from the Bombcell^61^ toolbox.

#### Noise units

were identified by unphysiological waveform characteristics: peak-to-trough duration < 0.05 ms or peak-to-trough duration > 15 ms, waveform baseline flatness > 0.7, peak-after-to-trough-ratio > 0.8, or exponential decay rate < 0.01 & exponential decay rate > 0.1

#### Non-somatic units

(axonal or dendritic) were identified by: peak-before-trough ratio > 3.0, peak-before-width < 0.15 ms, or peak-before-to-peak-after ratio > 3.0.

#### Multi-unit activity

comprised clusters failing single-unit isolation criteria: median amplitude < 30 µV, SNR < 2, noise cutoff > 5, total spikes < 100, refractory period contamination > 0.1, or presence ratio < 0.9

#### Well-isolated units

were all units that are not flagged as noise, non-somatic, or multi-unit active.

### Longitudinal Unit Tracking

To follow the same units across days, recordings from the last baseline day (Day −1), the acute day (Day 0), and 24 h later (Day 1) were concatenated with Supercat, preprocessed together (CatGT and DREDge motion correction), and spike-sorted as a single recording with Kilosort4, so that a cluster spanning the three sessions was treated as the same unit. Clusters were then curated as described above. Only well-isolated units present in all three sessions (presence ratio > 0.9; see above) were analyzed (n = 1692 units from 8 mice; Fig. 1D).

Validation of unit tracking. To test whether the identities followed the same neuron across days, unit properties were compared between recording windows, using nearby units as nulls. For each day, two adjacent 7.5 min blocks (A and B) were assigned from a period with no injection. On Day −1 and Day 0, these were the 15 min before the injection; on Day 1, it was the whole 15 min recording. Within each block, the mean waveform template (up to 500 randomly selected spikes, 15 channels nearest the peak), soma location (monopolar triangulation), and spatial decay constant of amplitude across channels were computed for every unit.

For a unit i in block x and candidate unit j in block y, six waveform metrics were computed:

1. the Pearson correlation between templates
2. normalized Euclidean distance between templates
3. log ratio of peak-to-peak amplitude
4. correlation between peak-to-peak amplitudes on each channel
5. log ratio of spatial decay
6. distance between soma locations.

Pairs in which either unit fired fewer than 50 spikes were excluded. Each metric was converted to a similarity: correlations were Fisher z-transformed, and distances and log ratios were sign-inverted absolute values. Similarities were averaged across all block pairings between days. To give a combined waveform score, each similarity was z-scored across all pairs within a comparison, and the six z-scores were averaged.

The candidates for each unit i were the unit itself and every other unit on the same probe whose soma lay within 100 µm. Rank-1 identification was the fraction of units more similar to themselves than every one of their neighbors. Units with no neighbor within 100 µm were excluded from this measure.

Comparisons between days were benchmarked against a within-day ceiling, the agreement expected for the same neuron in the absence of the recording gap. Firing statistics were not used, so that drug-induced changes in activity could not affect the tracking assessment.

### Cell-type Classification

Putative cell types were assigned using the CellExplorer^28^ cell-type classification pipeline, which we re-implemented in Python. Units flagged as non-somatic (see Unit Classification and Quality Metrics) were kept as a separate class. The remaining units were classified from waveform and firing-property features. Units with a trough-to-peak time ≤ 0.425 ms were classified as narrow interneurons, a group that tends to include parvalbumin-expressing cells^28^. Units with a trough-to-peak time > 0.425 ms and an autocorrelogram time rise constant τ*_rise_* > 6 *ms* were -classified as wide interneurons. All other units were classified as pyramidal cells, which typically have wide waveforms, fire in bursts, and have mean firing rates below 2 Hz.

### Drug Protocol

Psilocybin (#7437-001) was obtained from the National Institute on Drug Abuse Drug Supply Program. Psilocybin was dissolved in sterile saline to a concentration of 0.2 mg/mL and administered at a dose of 1 mg/kg (injection volume: 5 mL/kg). Psilocybin and saline were administered subcutaneously at equivalent injection volumes. After 7 days of handling and habituation to head-fixation (Days −11 to −4), each mouse received saline injections on three baseline days (Days −3 to −1) and psilocybin on “acute” day (Day 0), in that fixed order. The first two saline sessions also acclimated mice to the injection. Saline injections were always given before psilocybin given its long-term effects.

### Histology & Probe Track Reconstruction

Before implantation, probe shanks were coated with a lipophilic fluorescent dye (Vybrant™ DiI, 1 mg/mL in ethanol, Invitrogen). After recordings, mice were deeply anesthetized with isoflurane, and probes were explanted for reuse. Mice were then transcardially perfused with ice-cold phosphate-buffered saline (PBS, 10 mM) followed by 4% paraformaldehyde (PFA) in PBS. Extracted brains were post-fixed in 4% PFA for 24 h at 4°C and stored in PBS until sectioning.

Coronal sections (100 µm thick) were cut on a vibratome (Leica) and counterstained with DAPI. Sections were imaged using a widefield fluorescence microscope. Probe trajectories were reconstructed by registering images to the Allen Mouse Brain Common Coordinate Framework (CCFv3)^62^ using an adapted SHARP-Track^27^ pipeline. DiI fluorescence tracks were manually traced to map recording sites to anatomical subregions. For 2 mPFC probes and 1 vHPC probe without histology, recording sites were assigned to regions using the known implantation trajectories.

Region groupings. Recording sites were assigned to the regions analyzed using Allen CCFv3 acronyms: HPF (CA1, CA3, DG, SUB); MB (SNr); DORpm (LP, PIL, POL, SGN); and DORsm(MG, PP, PoT). Regions with fewer than 20 units in total were combined into “Other” (CP, LSr, RSP, TTd, ZI) and were not analyzed separately.

Note: The Fig 1C vHPC section is anterior to the targeted insertion plane (AP −2.92 mm) and was chosen to show the zona incerta, which are not visible in the coronal slice at that insertion plane.

### Classification of unit modulation

Single units tracked across the saline and psilocybin sessions were classified as activated, suppressed, or not modulated (n.s.) into two post-injection windows: an acute window and a persistent window. For each treatment, baseline firing rate distribution (mean and standard deviation) was built from 1 min bins in a 10 min pre-injection window (12–2 min before injection day). The firing rate in each post-injection window was the mean rate over the window. The acute window was 10-15 min after injection, and the persistent window was 5-10 min into the recording made approximately 24 h after the respective injection. Modulation was expressed as a z-score (post-injection firing rate - baseline mean firing rate) / baseline firing rate standard deviation, s.d.) Z-scores were clipped to ±10. Units were classified as activated (*z* > 2), suppressed (*z* < −2), or n.s. (|*z*| ≤ 2). Because the same units were recorded under both treatments, each unit received an acute and a persistent label under saline and psilocybin. Brain regions were assigned from histology and cell type from each unit’s mean waveform.

### Statistical Analysis

Paired comparisons of modulation categories. Within each effect (acute, persistent), saline and psilocybin were compared on units tracked across both treatment sessions. Marginal homogeneity of the three-category distribution (activated, non-modulated, suppressed) was assessed with the Stuart-Maxwell Test. Directional effects were tested with exact McNemar tests, comparing activated vs. not activated and suppressed vs. not suppressed. P-values were Bonferroni-corrected for the two directions. The same tests were run within each brain region and cell type that had at least 20 units. Analyses were performed with statsmodels.

Bayesian mixed-effects model. To account for the clustering of units within probes and animals, treatment effects on the modulation category were also estimated with a Bayesian categorical (multinomial logit) mixed-effects model. Modulation category was the outcome, with non-modulated (n.s.) as the reference category, and treatment was the fixed effect, with saline as the reference. Random intercepts were included for subject, probe nested within subject, and unit nested within probe. Models were fit in Bambi with Bambi’s default weakly informative priors, using the No-U-Turn sampler in PyMC, with 4 chains of 1,000 tuning and 2,000 posterior draws each (target acceptance 0.95). Convergence was confirmed by R^ < 1.01 and bulk effective sample size > 400 for all treatment parameters. Effects are reported as odds ratios (ORs; the exponentiated posterior mean log-odds) of psilocybin relative to saline, i.e., the change in the odds of a unit being activated or suppressed, rather than non-modulated. ORs are given with 95% credible intervals (CrIs), obtained by exponentiating the bounds of the 95% highest-density interval on the log-odds scale, and the probability of direction (pd), the posterior probability that an effect has the same sign as its posterior mean (range 0.5–1). An effect was considered credible when its 95% CrI excluded 1 (0 on the log-odds scale). Separate models were fit for the acute and persistent effects and, in per-group analyses, for each brain region and cell type.

Acute-to-persistent transitions. For each treatment, units classified in both the acute and persistent windows were cross-classified by acute and persistent response classes (activated, non-modulated, suppressed). Within each acute class, the distribution of persistent classes was compared between saline and psilocybin with a χ² test of independence (Bonferroni-corrected for the three acute classes). For acute classes with a significant χ² test (acute activated and acute non-modulated), each persistent class was then compared between treatments with a 2 × 2 Fisher’s exact test (that class vs. all others), Bonferroni-corrected for the six comparisons. Transitions were also modeled with the Bayesian categorical mixed-effects model described above, fit separately for each acute class on the units in that class. Persistent response class was the outcome (non-modulated as the reference), treatment was the fixed effect (saline as the reference), and random intercepts were included for subject, probe nested within subject, and unit nested within probe. Because non-modulated was the reference outcome, the model estimates the odds of persisting as activated or suppressed relative to non-modulated. Shifts toward the non-modulated class were therefore evaluated with Fisher’s exact tests.

Time courses. Spike counts were binned in 1 min bins, converted to firing rates and smoothed with a Gaussian kernel (σ = 2 bins). Smoothing was applied separately within each recording session so that it did not extend across session boundaries. Smoothed rates were then *z*-scored to each unit’s 15 min pre-injection baseline (mean and s.d. of the 15 baseline bins), and *z*-scores were clipped to ±10. Units with zero baseline variance were excluded. Traces show the mean ± s.d. across units in each response class. To match the duration of the saline sessions, psilocybin-day traces were restricted to 15 min before and 15 min after injection (Fig. 4A). For the persistent effect (Fig. 4C), Day 1 firing rates were z-scored to the acute-day (Day 0) pre-injection baseline.

Response magnitude. Response magnitude was quantified as |z|, where z was the z-score of a unit’s mean firing rate 10–15 min post-injection window relative to a 10 min pre-injection baseline taken from the center of the 15 min pre-injection period, and clipped to ±10. |z| was computed identically for saline (Day −1) and psilocybin (Day 0). |z| was compared between saline and psilocybin across the 1692 tracked units with a two-sided Wilcoxon signed-rank test on paired units, and per-animal medians were plotted as paired points. To compare the shape of the two |z| distributions, a Q-Q plot was generated to plot 1st-99th percentiles of psilocybin |z| against the same percentiles of saline. Percentiles were computed for each condition across all units. To test where the distributions differed, we computed the psilocybin – saline difference in |z| at each decile (10th – 90th). 95% confidence intervals for each difference came from a paired bootstrap over units (2,000 resamples, keeping each unit’s saline and psilocybin values together). A decile was considered significantly shifted if its 95% CI excluded zero.

Linear mixed-effects model. We modeled each unit’s Day 1 z-score as a function of its late acute response. *z_late_* was the mean z-score from 55 to 60 min post-injection on Day 0, taken from the z-score time course after Gaussian smoothing (σ = 2 min). *z_day_*_1_ was the z-score of the mean firing rate over 5-10 min of the Day 1 recording. Because a shared baseline could artificially inflate both z-scores together through baseline noise, *Z_late_* and *z_day_*_1_ were computed from separate, interleaved halves of the baseline (even and odd minutes). The model was fitted once for each assignment of the halves and with the full shared baseline. As the shared baseline minimally altered the results, that data was reported. The model was

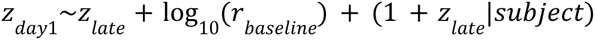

Baseline firing rate (log_10_ of the mean baseline rate, mean-centred) was included as a covariate to account for rate-dependent differences in z-score scale. Each animal had a random intercept and a random slope for *z_late_*, so the fixed-effect slope is the average across animals and its confidence interval incorporates between-animal variability in the slope. To test if initial response added any predictive power, *z_early_*, the mean z-score from 13-18 min post-injection on Day 0, was added to the model as well *z_day_*_1_ ∼*z_late_* + *z_early_* + log_10_ (*r _baseline_*) + (1 + *z_late_* |*subject*). Models were fitted by restricted maximum likelihood (REML) with statsmodels in Python. We report Wald 95% confidence intervals for the *z_late_* slope, which gives the change in Day 1 z-score per unit of late acute z-score. P-values are from two-sided Wald z-tests of the null hypothesis that the slope is zero.

Animal-level analysis. Because there were only eight animals, we also tested the effect without random effects. We fit the same fixed-effects model (Day 1 z-score on *z_late_* and log _10_ baseline rate) separately in each animal by ordinary least squares, and report the number of animals with a positive *z_late_* slope.

## Data and Code Availability

The data and code supporting this study will be made publicly available.

## Acknowledgments

We thank Jackson Freiman for assistance with psilocybin preparation and Adam Charles for general advice on data analysis. We also thank the NIDA Drug Supply Program for providing psilocybin. TD was supported by NIH/NIMH F31MH140536-01A1 and the Johns Hopkins University XDBio Fellowship. TD, AC, SK, AG, and TH were supported by NIH U01NS115587-04 (NIH BRAIN Initiative). TD, AG, KT, and TH were supported by a Johns Hopkins University Discovery Grant.

